# Neural mechanisms of resource allocation in working memory

**DOI:** 10.1101/2024.05.11.593695

**Authors:** Hsin-Hung Li, Thomas C. Sprague, Aspen H. Yoo, Wei Ji Ma, Clayton E. Curtis

**Affiliations:** Department of Psychology, New York University, New York, NY 10003, USA; Center for Neural Science, New York University, New York, NY 10003, USA; Department of Psychology, The Ohio State University, Columbus, OH 43201, USA; Department of Psychological and Brain Sciences, University of California, Santa Barbara, CA 93106, USA

## Abstract

To mitigate capacity limits of working memory, people allocate resources according to an item’s relevance. However, the neural mechanisms supporting such a critical operation remain unknown. Here, we developed computational neuroimaging methods to decode and demix neural responses associated with multiple items in working memory with different priorities. In striate and extrastriate cortex, the gain of neural responses tracked the priority of memoranda. Higher-priority memoranda were decoded with smaller error and lower uncertainty. Moreover, these neural differences predicted behavioral differences in memory prioritization. Remarkably, trialwise variability in the magnitude of delay activity in frontal cortex predicted differences in decoded precision between low and high-priority items in visual cortex. These results suggest a model in which feedback signals broadcast from frontal cortex sculpt the gain of memory representations in visual cortex according to behavioral relevance, thus, identifying a neural mechanism for resource allocation.

---

Working memory (WM) holds information temporarily and supports decision-making, learning and planning. A signature of WM is that it has severe limitations in its capacity; the precision of WM reports declines steeply as the number of items being stored increases ^1–6^. In addition, neural activity that supports WM is inherently noisy and the quality of WM representations as indexed by behavioral reports fluctuates across trials ^7–14^. Thus, decisions based on WM are always subject to uncertainty.

Whereas the total resource or capacity of WM is limited, individuals have some control over how their memory resource is allocated. Humans can prioritize and maintain more precise memory for items that are more relevant ^15–20^ or associated with higher reward ^20,21^. Moreover, people not only show smaller memory errors, but also report lower uncertainty about their memory for high-priority items ^17^, indicating that they have access to priority-dependent uncertainty when making WM-based decisions.

Allocating WM or attentional resource across spatial locations leads to corresponding modulations of neural responses in cortical retinotopic visual maps. In both perception- and WM-based tasks, attention boosts the neural responses at the prioritized locations ^18,22–33^. While these studies provided important insights into how the brain allocates WM or attentional resource flexibly, they primarily focused on neural responses aggregated over trials where items were maintained in WM at a specific priority level or prioritized at a specific attentional state (e.g., attended vs. unattended). Without quantifying the quality of WM on a trial-by-trial basis, previous studies can not address how prioritization affects uncertainty, nor did they investigate how resource allocation or prioritization could fluctuate across trials. In addition, without providing a linking model, how WM prioritization or attentional modulations observed in neural responses are related to behavior remain unresolved. Moreover, the cortical neural networks controlling the trial-by-trial allocation of WM resource remain unknown.

Theoretical ^34–40^ and experimental ^41–43^ work in the perceptual domain has demonstrated that sensory neurons can use probabilistic population codes to jointly encode stimulus features and their associated uncertainty. Recently, we extended this theory to spatial WM ^13^ and demonstrated that neural activation patterns in extrastriate visual and parietal cortex jointly encode the remembered spatial position and its trial-by-trial uncertainty. Building on these findings, here we test the hypothesis that behavioral relevance sculpts the gain of population neural response, and with the same computational principle—probabilistic population codes—human brains jointly represent the content and uncertainty of multiple memorized items with varying priority levels that are concurrently held in WM (Figure. 1C). We built a generative model-based decoder that used fMRI responses to demix and decode two items held in WM with different behavioral priorities. This critical extension of the probabilistic decoding approach ^13,41,43–45^, from one to multiple dimensions, allowed us to decode the content and uncertainty of multiple WM items on a trial-by-trial basis.

**Figure 1.**
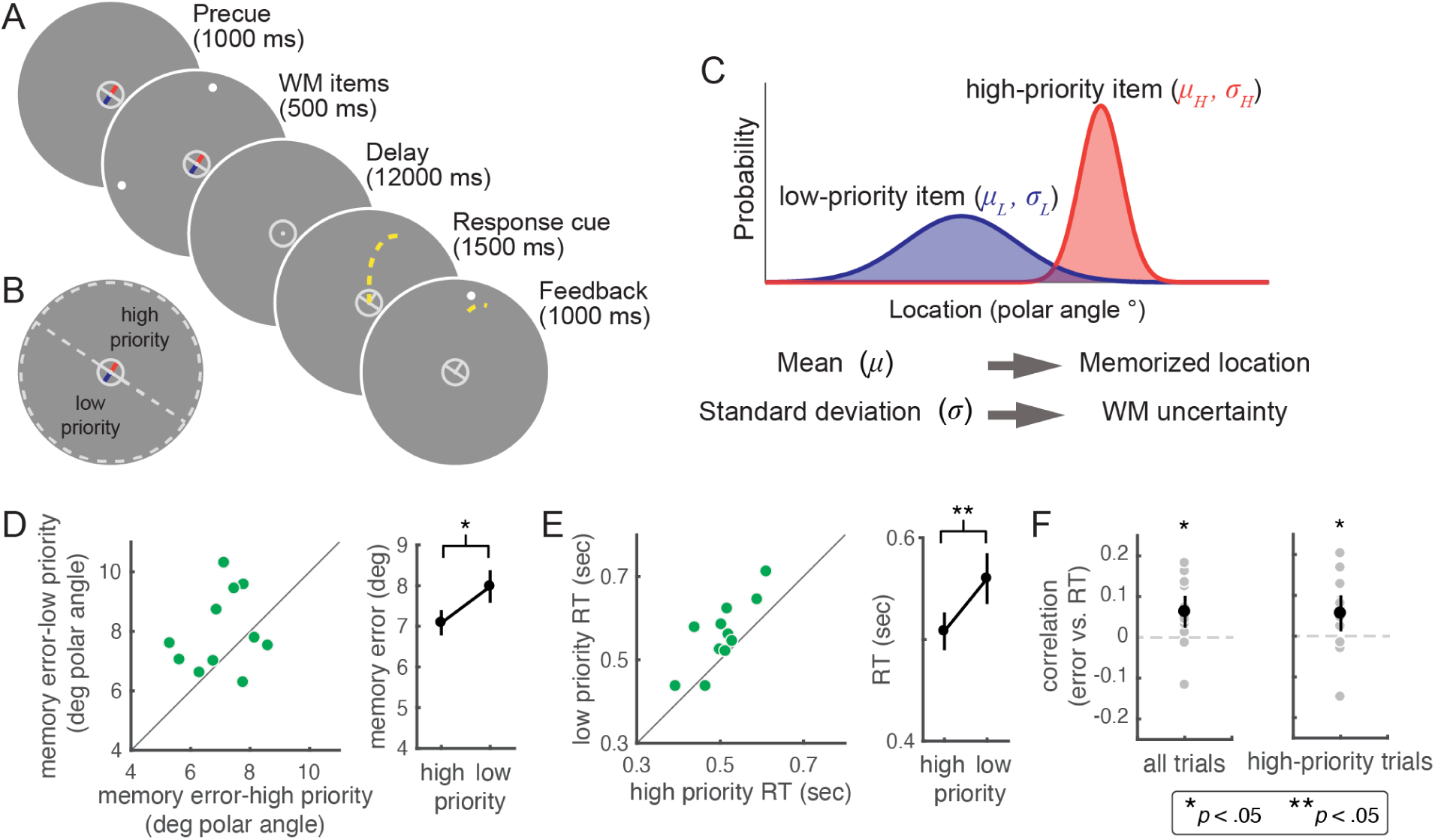
Procedures and behavioral performance. (A) Each trial started with a precue, which divided the circle into two segments (semicircles) by the orientation of a gray bar, randomly oriented on each trial. The color of each of two orthogonal bars in the precue also informed participants in which segment the high-priority item would appear. After the precue onset, two items simultaneously appeared, one in each segment. The location (polar angle) of each item was pseudo-randomly sampled within its semicircle. The WM items were followed by a 12-sec delay. At the end of the delay, a response cue prompted participants to make a memory-guided saccade to the memorized location of one of the items. The high-priority item was probed twice as frequently as the low-priority item The true target location was then re-presented as feedback. (B) Schematic illustrating the information conveyed by the precue. The dashed lines were not presented during the experiment, but depict how participants used the precue, as it divided the full circular aperture into two semicircles. The orientation of this division was random across trials. (C) According to probabilistic population coding theory, two items held in WM would be represented as two probability distributions, with means and standard deviations representing the memorized location and the associated uncertainty, respectively. Furthermore, high-priority items are represented with lower uncertainty. (D) Memory errors were smaller for high-compared to low-priority items (mean absolute error in degrees polar angle. Left: each dot represents one participant. Right: data points represent mean ± s.e.m. (E) Saccade response times are faster for high compared to low-priority items, plotted in the same format as in (D). (F) Memory errors and response times correlate. The black dots/lines represent mean ± s.e.m. Each gray dot represents one participant. Left, the correlation computed by all the trials; right, the correlation computed using only the trials that probed the high-priority item.

To preview, we measured fMRI responses while participants remembered the locations of two targets with different levels of behavioral relevance. As hypothesized, the gain of neural responses varied with target priorities. Higher priority targets were decoded with smaller error and lower uncertainty. These neural differences in prioritization predicted memory behavior as well. We also found evidence that trial-wise delay activity in frontal cortex predicted differences in decoded precision between low and high-priority items in visual cortex. Together, these results support a model in which activity in association cortex is the source of feedback signals that sculpt the gain of WM representations in visual cortex according to behavioral relevance.

## Results

We studied how people allocate WM resource during a memory-guided saccade task. In each trial, participants were presented with two items, each placed in one of two segments (semicircles) of the visual field that were divided by a precue. The precue identified the item priorities (Figure 1A and 1B). Participants were asked to remember the locations of the two items over a 12-second memory delay. Priority was manipulated by the probability that an item would be cued at the end of the trial. After the memory delay, a response cue appeared, prompting participants to report the location of one of the two items by a saccadic eye movement. Participants were twice as likely to be asked to make a memory-guided saccade to the high compared to the low-priority item. From previous psychophysical studies, we know that participants improve their performance by allocating more WM resource to the high-priority item ^17,20^. To identify target regions of interest (ROI) for decoding, we used population receptive field (pRF;^46^) modeling to identify visual field maps in visual cortex (V1, V2, V3, V3AB), parietal cortex (intraparietal sulcus IPS0, IPS1, IPS2, IPS3) and prefrontal cortex (superior precentral sulcus sPCS, inferior precentral sulcus iPCS).

### Priority affects behavioral performance in WM

At the behavioral level, the magnitude of the memory error was smaller for high-priority items than for low-priority items (permutation test, *p* = .043), indicating that participants maintained more precise memory for items that are more relevant for the task (Figure 1B). Prioritization was also reflected in the response time: saccade response time was shorter for high-priority than low-priority items (Figure 1D; permutation test, *p* < .001). In addition, we observed a correlation between saccade response time and the magnitude of memory error across all trials (*M* = .063, bootstrapping test, *p* = .011; Figure 1E) or among high-priority items (*M* = .056, bootstrapping test, *p* = 0.028; Figure 1F). Response time shows a robust correlation with confidence reports ^47^, and is often considered as an indicator of decision uncertainty or confidence ^48–50^. Thus, our results indicate that participants’ uncertainty reflects the precision of their memory.

### Bayesian encoding-and-decoding model for multiple items

By extending a previous model for one item ^13,41,44,45^, we built a generative encoding model for voxel activity patterns in response to two items with varying priority. The voxel activity pattern was modeled as a multivariate normal distribution. The mean of this distribution was determined by each voxel’s spatial (polar angle) tuning curve, which was modeled as a weighted sum of eight basis functions that evenly tiled visual (polar angle) space (Figure 2A). The model assumes that neural populations encode two items with different gains based on their priority levels. Thereby each voxel’s response to two simultaneously presented items was computed as a weighted sum of its response to each item presented individually (Figure 2B). Even though conceptually there are two gain factors, when both gain factors are allowed to vary, at least one of the two factors become interchangeable with the scale of the voxel tuning function (Figure 2A). Therefore, we fixed the weight (gain) of the high-priority item at 1 and fit the weight of the low-priority item as a free parameter. The covariance of the distribution was estimated using the empirical noise covariance and a theoretically-driven covariance matrix, as in previous implementations (see Methods) ^13,44,45^.

**Figure 2.**
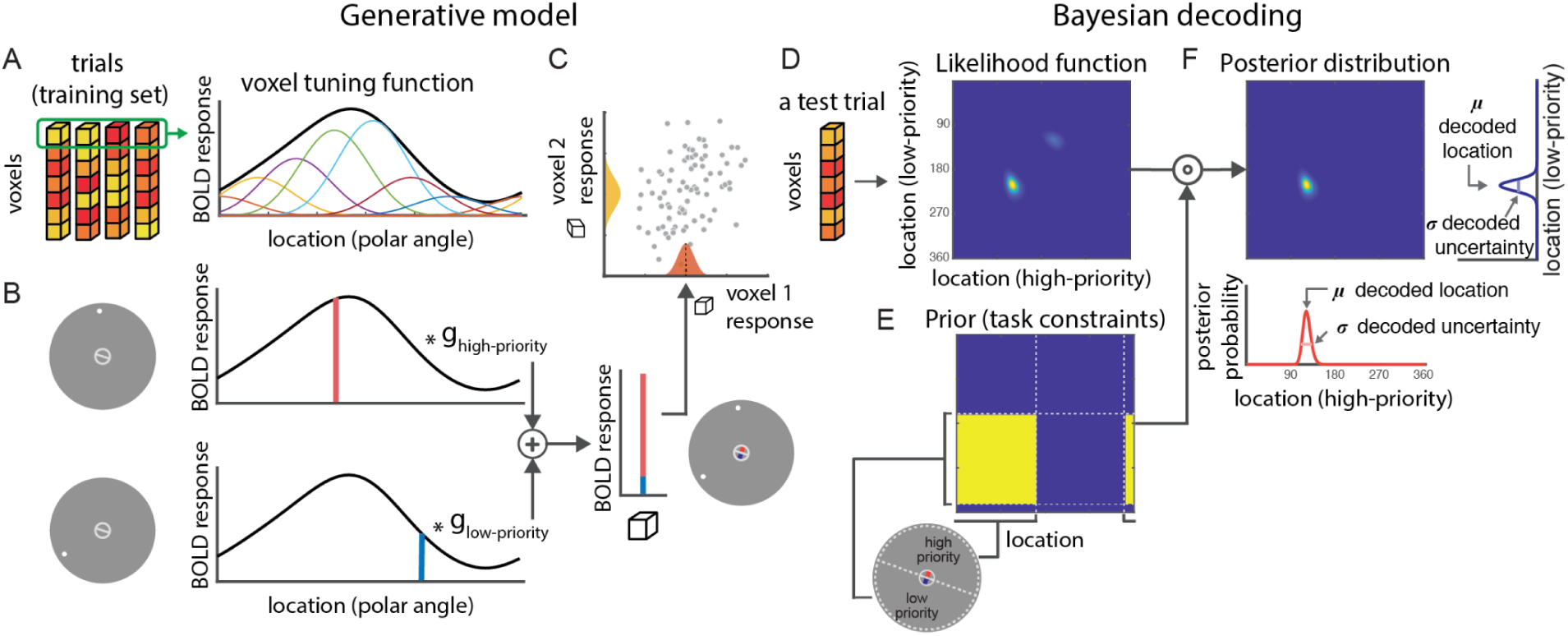
Generative model-based decoding analysis. (A-C) The generative model (encoding model). (A) Using trials in the training set, each voxel’s polar angle tuning function (the thick black curve) was modeled as a weighted sum of eight basis functions (the thin colored curves) evenly spaced across polar angle (0° to 360°). (B) A voxel’s mean response to two items with different priority levels were modeled as a weighted (by two gain factors g_high-priority_ and g_low-priority_) sum of the voxel’s response to each of the items presented alone. In practice, we fixed the gain of the high-priority item (g_high-priority_) at 1 and fitted the gain of the low-priority item (g_low-priority_) as a free parameter using all trials in the training set. (C) Voxel activity pattern was modeled as a multivariate normal distribution. The scatter plot is a schematic of a bivariate normal distribution representing the response of two hypothetical voxels (voxel 1 and voxel 2), where the black dashed line represents the mean response of voxel 1. The mean of the distribution was determined by the mean response of each voxel. The covariance of the distribution was estimated using the empirical noise covariance and a theoretically-driven covariance matrix (see Methods). For each ROI, the free parameters in the generative models were estimated using leave-one-run-out cross-validation. (D-F) Bayesian decoding model. (D) Given the voxel activity pattern of a trial in the test set (held-out run), a two-dimensional likelihood function was computed, where the horizontal and vertical dimensions represent the location of high and low-priority items (polar angle) associated with g_high-priority_ and g_low-priority_ respectively. Note that the likelihood function would only be a symmetric matrix if the estimated weights for the high- and low-priority items are the same. (E) The decoder further utilizes the knowledge given by the precue, including how the visual field is divided into two semicircles (each covers a range of 180°) and the associations between the semicircles and their priority levels. This information was implemented as the prior, a task-constraint matrix consisting of 1s and 0s. (F) A two-dimensional posterior distribution was computed by pointwise multiplication between the likelihood function and the prior. We computed one-dimensional posterior distributions for the high- and low-priority items by marginalizing over the two-dimensional posterior distribution. For each item, the circular mean and circular standard deviation of the decoded posterior distribution represent decoded location and decoded uncertainty, respectively.

We next assumed that a decision-maker infers WM content using the knowledge of the generative model. Thus, we implemented a Bayesian decoder that ‘inverted’ the generative model (Figure 2D-2F). While previous studies utilized Bayesian decoders for a single stimulus feature ^13,41,44,45^, here, we faced the additional challenge of demixing two items from the response in the same neural population. To achieve that, we implemented a decoder that understood which hemifield was prioritized according to the precue (Figure 2E). On each trial, we decoded the location of each of the two items as a probability distribution over all possible locations within its segment. We then used the circular mean of the probability distribution to represent the decoded location, and the standard deviation of the distribution to represent decoded uncertainty.

### Priority sculpts WM content and uncertainty held in neural populations

We first examined whether priority affects the accuracy of WM content readout from neural populations. In an example participant, we visualized decoded location as a function of actual item location (top rows in Figure 3A and 3B), and as the distribution of decoding errors for each brain region of interest (ROI; bottom rows in Figure 3A and 3B). In all ROIs, the distribution of decoding errors showed a single peak around 0°, indicating decodable WM content. Moreover, we observed that the decoding error was smaller for the high-priority than the low-priority item, mirroring the effect of prioritization at the behavioral level. At the group level (Figure 4A), we found that all the ROIs showed above-chance decoding by comparing group-averaged decoding error to the null distributions estimated by permutations (dashed lines in Figure 4A; permutation test *p* < 0.05 for all the ROIs). An ANOVA on the magnitude of decoding error showed a significant main effect of ROI (permuted two-way repeated-measures ANOVA, *F*(9,90) = 36.13, *p* < 0.001; Figure 4A) and an interaction between ROI and priority (*F*(9,90) = 5.01, *p* < 0.001). Testing the effect of priority on individual ROIs, we found that decoding error was significantly lower for the high-priority than the low-priority item in V3AB and IPS0 (permutation test *p* < 0.05; unless otherwise noted, we report *p* values corrected for multiple comparisons across ROIs via false discovery rate [FDR] with *q* = 0.05; Figure 4A). We observed similar results when using the variability of decoded locations as an index for the decoder’s performance (Supplementary Figure 1).

**Figure 3.**
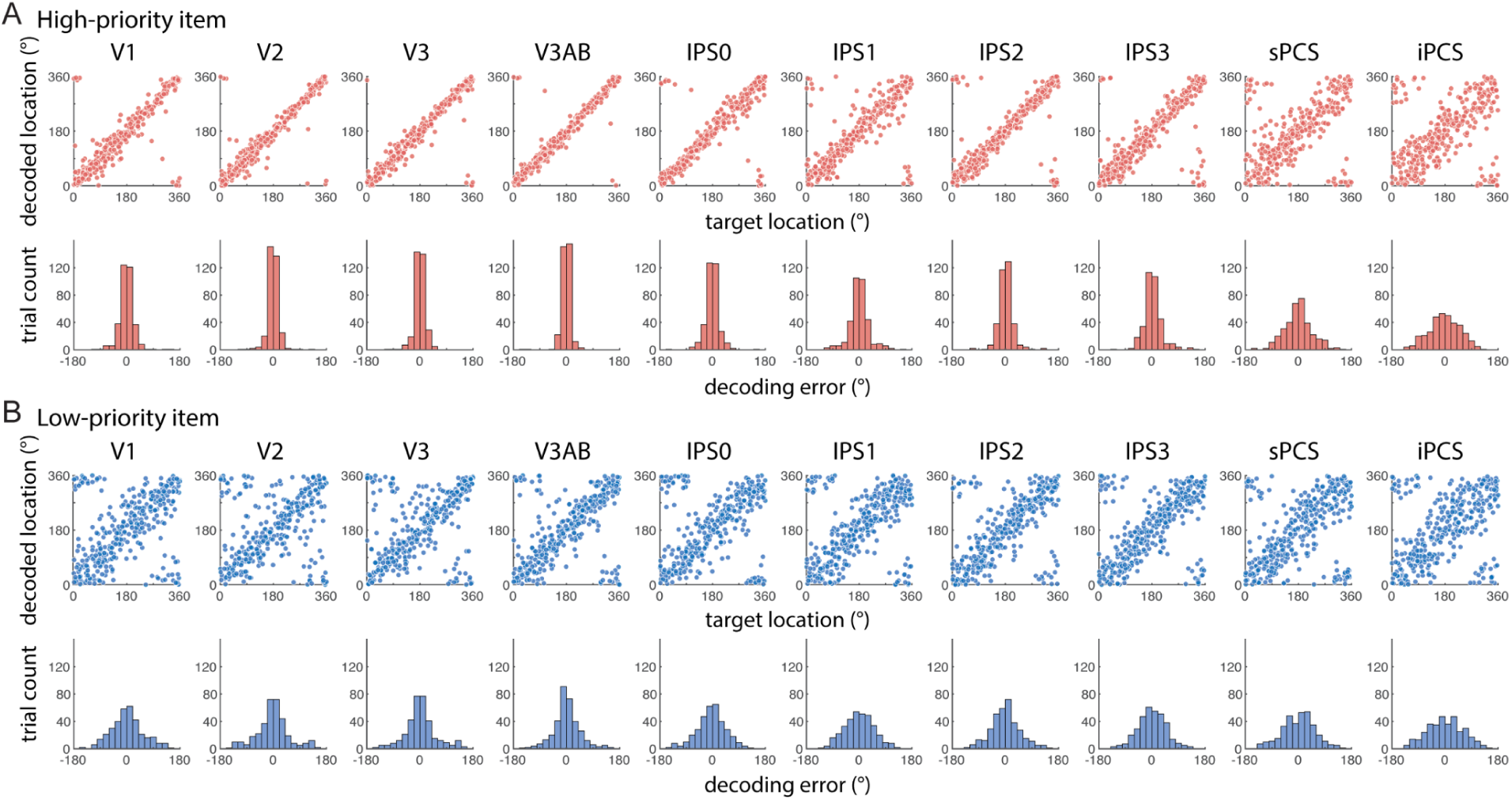
Decoding 2 items at a single-trial level from an example participant. (A) Decoding results for the high-priority items. Top: Decoded location (y-axis) plotted against actual target location (x-axis). Bottom: The distribution of decoding error, computed as the decoded location minus the actual target location. (B) Same as (A) but for low-priority items.

**Figure 4.**
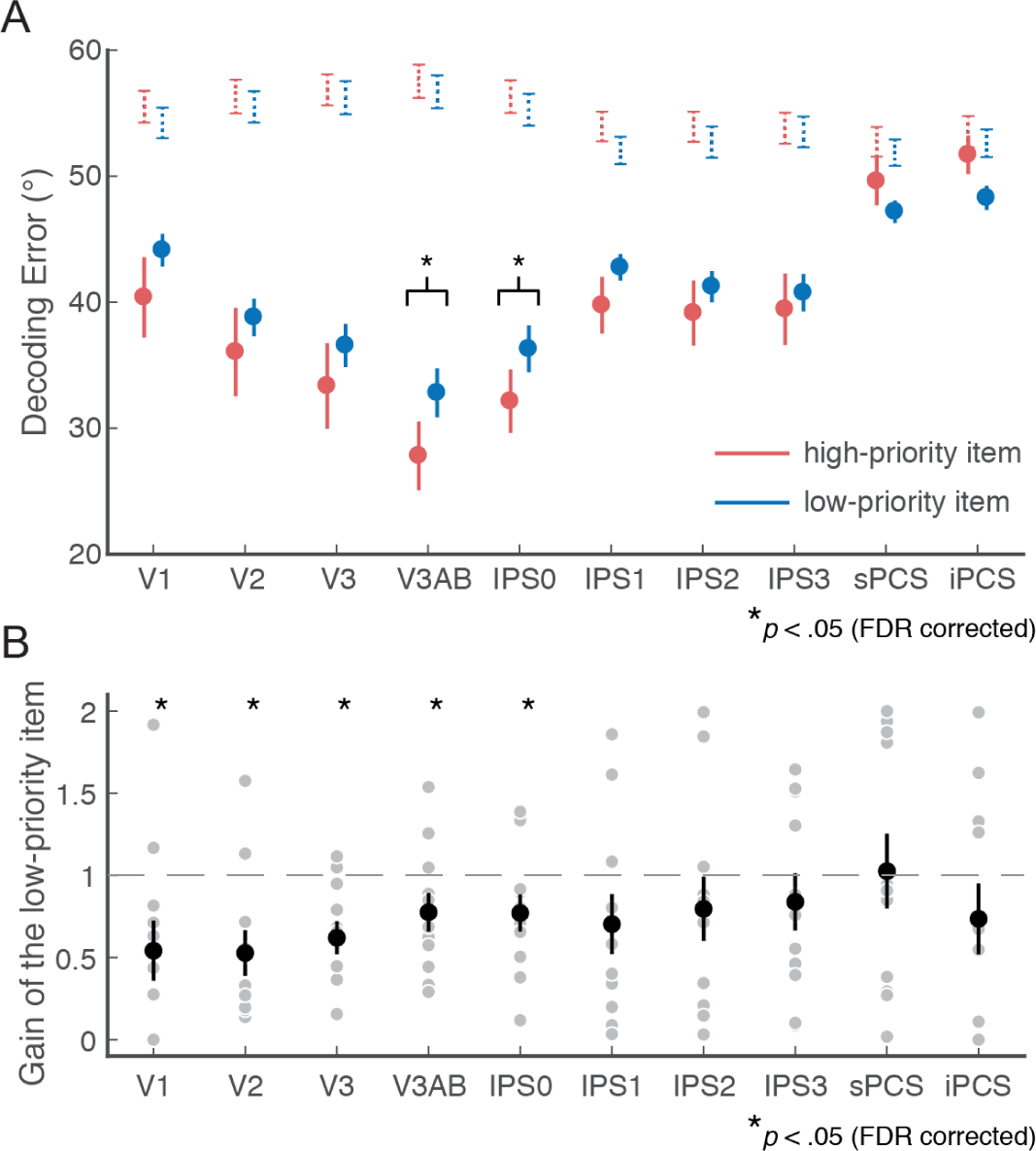
Decoding error and the estimated gain factor reflect behavioral prioritization. (A) Averaged magnitude (absolute value) of decoding error for the high- and low-priority items across ROIs. Data points represent mean ± s.e.m. Dotted lines represent 95% interval of the null distribution of decoding errors, estimated by randomly permuting the decoded likelihood functions across trials. (B) The estimated gain for the low-priority item. Black data points represent mean ± s.e.m. Gray data points represent individual participants. Asterisk symbols indicate bootstrapping tests against one. Gain for high-priority item was fixed at 1 (horizontal dashed line) (C) Behavioral prioritization (the magnitude of memory error of the low-priority item minus that of the high-priority item) plotted against the estimated gain for the low-priority item.

We found that priority impacted the neural gain allocated to the items held in WM. While we fixed the gain of high-priority items to be at 1, the estimated gain factor for low-priority items was smaller in visual cortex (V1-V3; V3AB) and IPS0 (bootstrapping test against gain = 1, *p* < 0.05; Figure 4B). In multiple brain regions, this gain factor closely tracked behavioral prioritization. Participants with a higher estimated gain for the low-priority item showed less behavioral prioritization, quantified as the magnitude of memory error of the low-priority item minus that of the high-priority item (Figure 5). Therefore, differences in the degree to which people prioritized the items can be explained by a mechanism in which the gain of an encoded item was modulated according to its priority.

**Figure 5.**
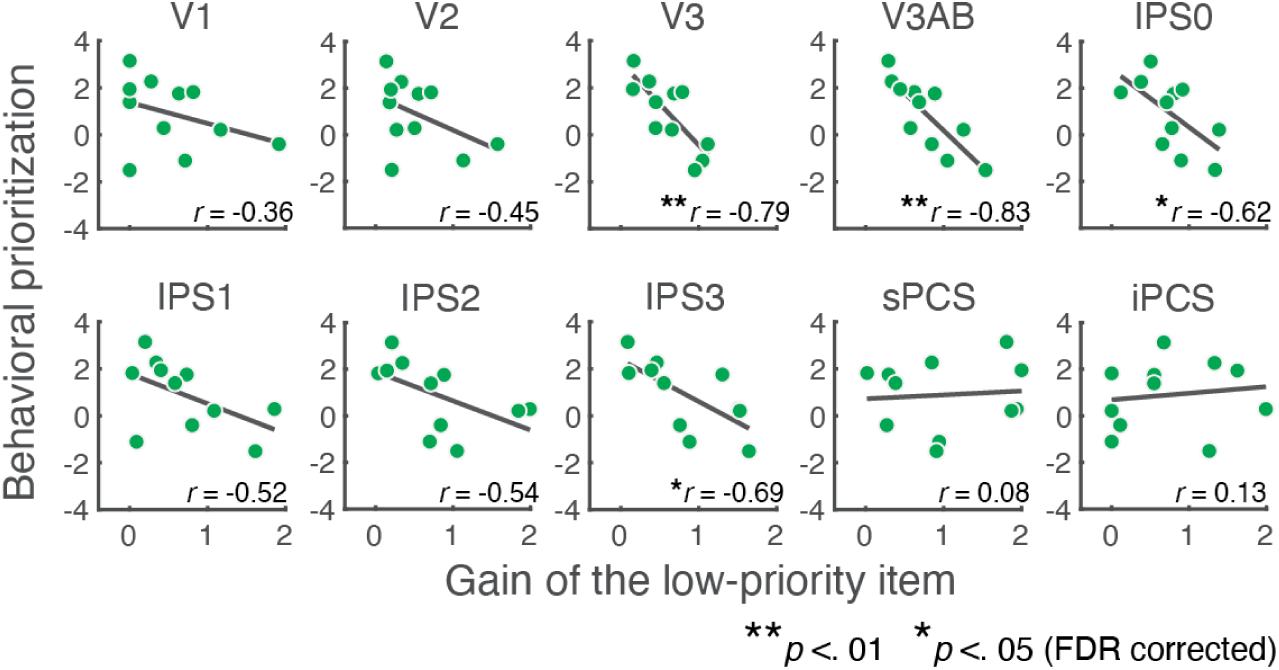
Gain modulations track individual differences in prioritization. Behavioral prioritization (the magnitude of memory error of the low-priority item minus that of the high-priority item) plotted against the estimated gain for the low-priority item.

People not only maintained more precise memory, but also reported being more certain about their memory for items with higher priority ^11,17^. Because neural populations utilize probabilistic information to represent the content held in WM ^13^, we hypothesized that priority would impact decoded neural uncertainty. Consistent with this hypothesis, we found that decoded uncertainty was lower for high-compared to low-priority items (permuted two-way repeated-measures ANOVA, main effect of priority, *F*(1,90) = 23.4, *p* < 0.01; Figure 6A). To further investigate whether decoded uncertainty reflected priority-dependent quality of WM on a trial-by-trial basis, for each trial, we estimated a WM prioritization index computed as the difference in decoded precision (the inverse of the square of decoded uncertainty) divided by the sum of decoded precision over the two items (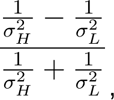 where *σ_H_* and *σ_L_* are the decoded uncertainty of the high- and low-priority item respectively). Conceptually, this index is similar to the ‘attentional modulation index’ often used in neurophysiological studies ^51–53^. Because we only had one behavioral measure per trial, we binned trials into quartiles of neural WM prioritization index (for each ROI), and computed each bin’s behavioral prioritization as the difference in the magnitudes of behavioral memory errors between trials in which the high- and low-priority items were cued. Remarkably, behavioral prioritization increased with neural WM prioritization in V1, V2 and V3AB (permutation test p < 0.05; Figure 6B). Therefore, larger differences in decoded uncertainty between the two items during the memory delay predicted stronger prioritization of memory-guided behaviors.

**Figure 6.**
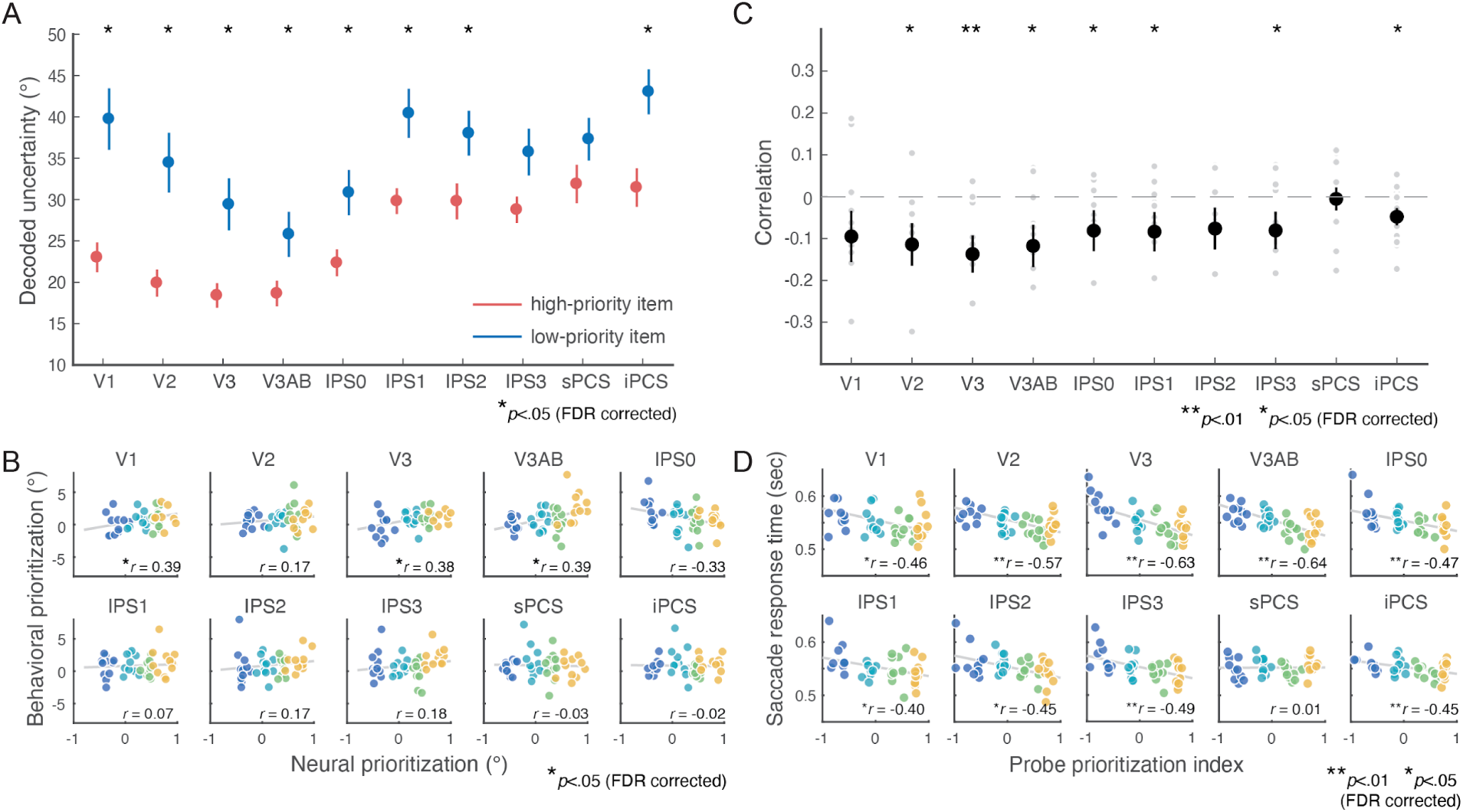
Decoded uncertainty reflects behavioral prioritization. (A) Decoded uncertainty for high- and low-priority items across ROIs. Data points represent mean ± s.e.m (B) Behavioral prioritization plotted against WM prioritization. WM prioritization was computed as the difference of decoding precision (inverse of the square of decoded uncertainty) divided by the sum of decoding precision over the two items. Each participant’s trials were binned into four bins with increasing WM prioritization (represented by four colors). For each bin, behavioral prioritization was computed as the magnitude of memory error for the low-priority item minus that of the high-priority item. (C) Correlation between saccade response time and the difference in decoded uncertainty between the item probed by the response cue and the unprobed item. Black data points represent mean ± s.e.m. Gray data points represent individual participants. (D) Visualizing the correlations in (C) in the format similar to (B). Each participant’s trials were ordered and binned into four bins (represented by four colors) based on the difference in decoded uncertainty between the probed and the unprobed item (x-axis) and the saccade response time was averaged across trials in each bin (y-axis).

Next, we tested whether the uncertainty of the neural representation predicted behavioral aspects of uncertainty. We leveraged findings from previous studies on perceptual or WM-guided decisions that response time correlates both with uncertainty and (negatively) with explicit reports of confidence ^47,49^. Thus, we treated the response time of memory-guided saccades as an implicit behavioral index of WM uncertainty, and asked if decoded uncertainty predicted response time. Indeed, we observed negative correlations between saccade response time and probe prioritization index (computed similar to WM prioritization index, except that the denominator was computed as the decoding precision of the probed item minus that of the non-probed item; Figure 6C and 6D). Specifically, when neural uncertainty was lower for the probed compared to the non-probed item, response times were faster. We observed similar results when correlating response time with decoded uncertainty of the probed item solely (Supplementary Figure 2). These findings further support a connection between the uncertainty represented by the neural population and the uncertainty at the behavioral level.

### Cortical network underlying control of WM resource

Our results so far showed that the quality of WM content represented by neural populations in visual cortex correlates with behavioral priority. To fully understand the neural circuitry involved in the control of WM resource, we tested the hypothesis that WM content represented in visual cortex is modulated by neural activity in brain regions that provide top-down signals associated with the allocation of WM resource.

We used the decoded uncertainty associated with each item on each trial measured from V3AB to represent the quality of WM content. We focused on V3AB because it exhibited the highest decoding performance (Figure 4A) and correlations with behaviors (Figure 5 and 6), a finding that is consistent with previous results when decoding a single item in spatial VWM ^13^. The ability to decode two items concurrently held in WM provided a unique opportunity to investigate two types of WM control. First, we quantified the total amount of allocated WM resource as the sum of decoded precision across the two items in each trial (Figure 7A). Second, we quantified how the WM resource was allocated to each item according to priority. As described in the previous section, this WM prioritization index was computed as the difference between the precision of the high- and low-priority items divided by the sum of decoding precision over the two items (Figure 7A).

**Figure 7.**
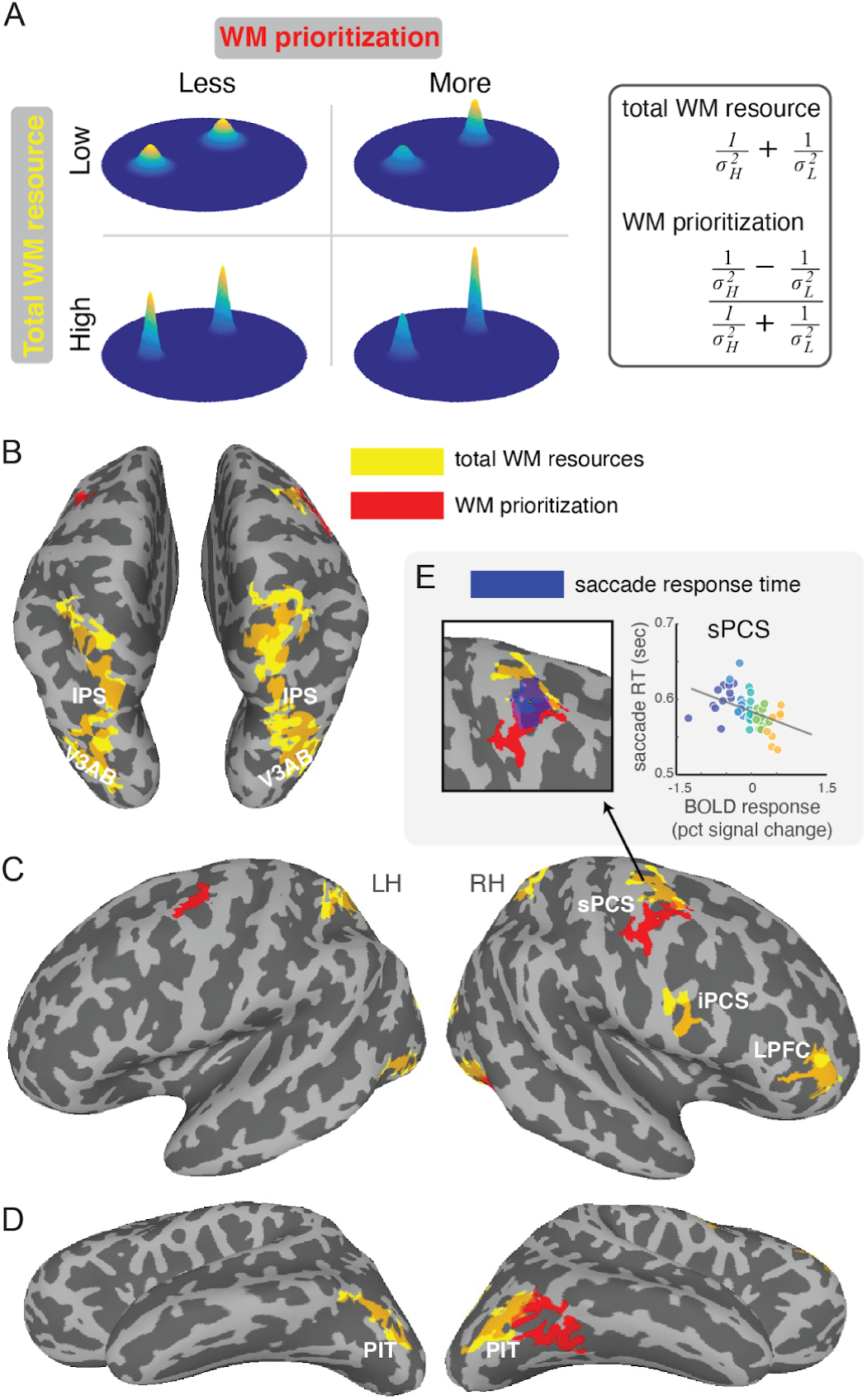
Cortical regions with responses fluctuating with the quality of WM. (A) We quantified total WM resource and WM prioritization using the decoded uncertainty from each trial. The schematic here illustrates how total WM resources, or WM prioritization, may vary while the other variable is held constant. In (B) to (D), clusters on the cortical surface exhibiting responses fluctuating with total WM resource and WM prioritization are labeled by yellow and red colors respectively. Here, the total WM resource and WM prioritization are computed using the decoded uncertainty from V3AB. For total WM resource (yellow), we identified significant clusters in (B) bilateral parietal cortex starting from V3AB and covering almost the entire IPS (intraparietal sulcus), several regions in (C) prefrontal cortex including sPCS (superior precentral sulcus), iPCS (inferior precentral sulcus) and LPFC (lateral prefrontal cortex), and bilateral clusters in (D) PIT (posterior inferior temporal cortex). For WM prioritization (red), the identified clusters were limited to (C) bilateral sPCS and (D) right PIT. (E) Delay period activity in right sPCS correlated with the response times of memory-guided saccades. Scatterplot shown for visualization, with trials binned by delay period response quartile per participant.

We postulated that the brain regions controlling the allocation of WM resource would exhibit BOLD amplitudes that fluctuates with either the total WM resource or WM prioritization index across trials. We identified these regions by whole-brain general linear (GLM) analyses, in which the regressor of interest was positioned at the time window of WM delay, and with amplitude varying trial-by-trial based on the total WM resource or WM prioritization. Significant clusters were identified by a voxel-level threshold (two-tailed t-test on the regression coefficient thresholding at *p* < 0.05), followed by a cluster-level size threshold (*p* < 0.05; see Methods).

Total WM resource decoded from V3AB covaried with delay period amplitude in the following regions (yellow clusters in Figure 7): (1) bilateral IPS (the posterior end of this cluster was in V3AB; Figure 7B) (2) right sPCS (3) right iPCS and (4) right lateral prefrontal cortex (rLPFC; the activated region occupied the anterior end of middle frontal gyrus MFG and inferior frontal sulcus IFS; Figure 7C) (5) bilateral PIT (posterior inferior temporal cortex; Figure 7D), a region positioned between the typical lateral and ventral visual field map clusters (see Supplementary Figure 3).

In a separate GLM, the prioritization of WM resource decoded from V3AB covaried with delay period amplitude in (1) right PIT and (2) bilateral sPCS (red clusters in Figure 7). No clusters with activity anti-correlating with total WM resource or neural prioritization were found. As we considered saccade response time as a behavioral correlate of WM uncertainty, we also conducted a GLM investigating the brain region with activity varying with saccade response time. We found a cluster at right sPCS with activity decreasing with saccade response time (Figure 7E). This cluster overlapped with the sPCS clusters that are associated with total WM resource and WM prioritization. Overall, we found that the control of WM resource recruits a large cortical network spanning parietal, prefrontal and temporal lobes.

## Discussion

We tested the hypothesis that human cortex represents the uncertainty of WM using a probabilistic neural code whose gain is modulated by the behavioral relevance of memoranda. In order to directly test this hypothesis, we developed a Bayesian decoding model to simultaneously estimate and demix the neural responses of two memory targets, as well as the impact of priority on those responses (Figure 2). These modeling innovations enabled us to demonstrate that the gain of neural activity in striate and extrastriate cortex tracked the priorities of the memoranda (Figure 4B). The neural gain of high-priority items was higher than that of low-priority items, and this difference predicted the behavior of how participants allocated their WM resource between the two items (Figure 5). Second, by leveraging a Bayesian decoder that used knowledge from the generative encoding model, we showed that items with higher priority were decoded with less error (Figure 4A) and less uncertainty (Figure 6A). Finally, the trial-wise amplitude of delay period activity in higher-order parts of prefrontal, parietal, and temporal cortex predicted the total amount of WM resource allocated to the two items, based on the total decoded uncertainty of the items in visual cortex. Moreover, delay period in prefrontal cortex predicted the degree with which the high-priority item was prioritized, here based on the difference in decoded uncertainty of the two items in visual cortex (Figure 7). Below, we discuss the implications of these associations between parameters of decoded WM representations and behavioral, neural, and theoretical variables.

The theory of probabilistic population codes explains how precise stimulus information can be read out from a population of noisy sensory neurons by positing that the population activity encodes a probability distribution over the stimulus feature ^34–40^. An advantage of probabilistic coding schemes is that they represent not only an estimate of the stimulus, but also uncertainty, whose use may improve decision-making ^40,54,55^. Indeed, stimulus uncertainty decoded from neural populations appears to be used by subjects performing perceptual decision making tasks^41–43,45^. In the case of working memory, we recently demonstrated that the content and uncertainty of memory could be decoded from visual cortex, and subjective reports indicated that participants were both aware of and used these representations of memory confidence ^13^. Building on this prior work and motivated by the probabilistic population coding framework, here, we manipulated how memory resource was allocated to two differently prioritized targets as a means to experimentally control changes in neural gain and investigate the impact of such changes on WM uncertainty represented by neural populations.

When quantifying the relative gain of each target in fMRI activation patterns during our generative modeling approach, we found that targets with lower priority were encoded with smaller gain by neural populations, and that differences in gain explained how individuals distributed their WM resource across the two memoranda (Figure 5). We assume that prioritization, the process of allocating WM resource, is similar to allocating attention weights. Thus, our results provide important extensions to previous reports that attention boosts neural responses at the attended location during perception ^22–29,33,56–58^. They also agree in general with computational models that use gain modulation to explain attentional effects on visual perception ^59^ and errors in WM ^19^. During WM delays, the amplitude of fMRI activity in visual cortex scales with the priority of items stored in memory ^18,60^. Our findings here provide more direct evidence that the gain of the prioritized location was enhanced in visual cortex during WM, and moreover, this gain predicted the behavioral effect of this prioritization (Figure 5).

While the relationship between the gain of a neural population response and stimulus certainty is the central construct in probabilistic population coding theories ^40,61^, a direct test of this relationship has proven challenging. Previous studies estimated the BOLD response gains of differently prioritized items in WM ^18,57^, but they lacked a model to link gain modulations with changes in neural uncertainty and behaviors. Conversely, previous studies that relate decoded uncertainty, of a single item, to behaviors in perception ^41,45^ and WM ^13^ did not investigate the effect of gain changes on neural uncertainty. We directly addressed this problem by developing an encoding-and-decoding model that allowed us to demix the neural representations of multiple WM items whose gains were experimentally under control. Our model allowed us to estimate, and thus connect, gain and uncertainty. We found that increased gain of high-priority items impacted both the accuracy and uncertainty of decoded WM representations (Figures 4A and 6A) in a manner predicted by theories of probabilistic population coding ^40,61^. These findings reveal a neural basis by which people incorporate state- or attentional-dependent uncertainty when making decisions ^7,17,62^. Critically, nonspecific factors like arousal cannot explain the specific differences in gain and uncertainty we decoded between the high and low-priority items. Instead, they were unambiguously driven by the differential allocation of WM resource between the two items.

All of the decoded measures of WM quality that tracked target priority were strongest in visual cortex and were absent in frontal cortex (eg, gain in Figure 4B). Given that the frontal cortex typically shows robust persistent activity during WM ^28,63–65^, what role then might the frontal cortex play in WM? To address this question, we performed two analyses to test the hypothesis that persistent activity in frontal cortex, and perhaps other areas, reflects feedback signals associated with the allocation of limited WM resource. These feedback signals presumably target areas in visual cortex that store WM representations.

First, we identified cortical areas whose trial-wise amplitudes of delay period activity covaried with the total amount of WM resource allocated to both items, defined as the sum of the decoded precision in visual cortex across the two items (Figure 7, yellow). This metric bears resemblance to the total WM resource in models that posit that the neural mechanisms that support storage of items in WM are limited by a memory resource that can be flexibly distributed across items ^1,3^. Total resource decoded from visual cortex was predicted by the amplitude of delay period activity in cortical areas that match the frontoparietal attention network or dorsal attention network ^66–70^. Total resource was also correlated with delay period activity in the right lateral PFC, similar to a previous study in which we found that variability in delay period activity in this area predicted decoding accuracy in a single item WM task ^31^. Overall, the frontoparietal network we identified resembles cortical networks identified in many visuospatial WM tasks ^64,71,72^. Nonetheless, since our total WM resource metric includes the decoded precision of both high- and low-priority items, it might reflect non-specific fluctuations of cognitive factors like general arousal ^73–76^ or mental effort ^77–80^.

Second, we identified cortical areas whose trial-wise amplitudes of delay period activity covaried with the relative amount of WM resource allocated to low- and high-priority targets. By taking the difference, our WM prioritization index eliminated obvious non-specific nuisance factors. Remarkably, delay period activity in the sPCS bilaterally predicted WM prioritization in visual cortex (Figure 7, red). The human sPCS likely contains the macaque frontal eye field ^46,81^, which is the source of feedback signals that target and alter the activity neurons in visual cortex ^82–84^. Our results converge well with a recent study that applied TMS pulses during memory delay, and found that disruptions of sPCS activity during the memory delay diminished the prioritization effect on behavioral memory error ^85^. Therefore, feedback signals originating in human sPCS may sculpt the neural representations of WM targets stored in visual cortex according to the item’s priority.

Unexpectedly, we found clusters of voxels in area PIT whose delay period activity correlated with both total WM resource and WM prioritization (Figure 7D). Although not previously identified as a region which plays a critical role in WM, this area of temporal cortex has been recently linked to the control of attention. Recent findings from electrophysiology in macaques ^86,87^ and fMRI in humans ^70^ suggest that the posterior-and-inferior part of the temporal cortex, including PIT, is involved in controlling visual attention. Moreover, along with evidence from network-level functional connectivity studies ^88^, there are strong white matter tracts that connect the dorsal IPS - which we have previously linked to trial-level behavioral judgments of WM uncertainty ^13^ - to posterior temporal cortex ^70,89^ providing a pathway by which attention signals in dorsal and ventral streams may be coordinated, extending the attention network to the ventral temporal cortex ^90^. The PIT area we identified sits between two clusters of visual maps (lateral maps LO1 and LO2 and ventral maps hV4, VO1 and VO2; Supplementary Figure 3), and consistent with previous fMRI studies ^67,88,91^ is not retinotopically organized. We highlight that the anatomical location of this region nearby to multiple extrastriate retinotopic maps and directly connected with posterior parietal cortex via a white-matter tract may enable efficient modulation of feature-specific representations in neighboring and connected regions.

Critically, our results are enabled by our new analysis approach which demixed multiple simultaneous and overlapping neural representations within activation patterns. This represents a critical extension of model-based decoding approaches as previous studies only considered decoding for a single item ^13,41,43,45,61^. By further extending the generative model to account for additional features, objects, and their interactions, future studies can build on the same approach and investigate the role of probabilistic neural codes for joint representations of multiple features or for natural stimuli.

In summary, we extended probabilistic population code to explain the representations of WM content and uncertainty for multiple items with different priority levels. Moreover, feedback signals from frontal cortex control the allocation of WM resource across items by modulating the gains with which different items are encoded. It then follows that behaviorally, memory uncertainty stems from decoding or readout of these representations with varying gains .

## Methods

### Participants

Eleven participants (5 females) took part in the experiment. All participants had normal or corrected-to-normal vision. The experiments were conducted with the written, informed consent of each participant. The University Committee on Activities involving Human Subjects at New York University approved the protocols of the study. Participants received monetary compensation of $30 per hour.

### Task

Participants performed a memory-guided saccade task in the fMRI scanner. Each trial started with a precue presented at the center of the screen. The precue was a gray annulus divided into two semicircles by a gray line (a divider) presented at the center of the screen (Figure 1A). The orientation of the divider was randomly chosen for each trial. A dark gray aperture with a radius of 15° was presented on the screen throughout the experiment. Participants were instructed to imagine that the divider in each trial cut the entire aperture into two segments (semicircles; Figure 1B). In addition, the precue contained a set of priority cues, a blue and a red line, orthogonal to the divider. The colors of the priority cues informed the participants which segment will hold the high-priority item. The associations between the priority levels and the colors were randomly assigned for each participant. One second after the precue onset, two WM items (one high-priority and one low-priority item, each with a width of 0.65° and a duration of 500 ms) were presented concurrently on the screen with an eccentricity of 12° from the fixation point (the precue). The polar angle of the WM items were chosen pseudo-randomly so that each segment contained one WM item, and the WM items were at least 10 degree away from the orientation of the divider in terms of polar angle. The WM items were followed by a delay with a duration of 12 seconds, during which participants maintained their gaze at the fixation point while remembering the locations of both of the WM items. At the end of the delay, a response cue appeared. The response cue contained the original divider, and a gray line pointing at one of the two segments, prompting the participants to report the location of one of the two items by saccadic eye movements. In 67% of the trials, the high-priority item was probed by the response cue while the low-priority item was probed in the remaining trials.

### Setup

During the experiment, the visual stimuli were presented by an LCD projector located behind the scanner bore. Participants viewed the stimuli through an angled mirror with a field of view of 52° by 31°.

### Eyetracking

Gaze position was tracked throughout the experiment by EyeLink 1000 Plus infrared video-based eye tracker (SR Research) mounted beneath the screen inside the scanner bore operating at 500 Hz. A 13-point calibration routine was conducted at the beginning of each session and was repeated between runs when necessary.

### Behavioral data analysis

Gaze position was analyzed offline, and was used as our measurement of VWM report. We preprocessed raw gaze data using fully-automated procedures implemented within iEye_ts (github.com/tommysprague/iEye_ts) to remove blinks, adjust for drift over the course of a run, recalibrate gaze data trial-by-trial, automatically identify memory-guided saccades, and flag trials for rejection for behavioral analyses. The details of the parameters used in preprocessing the gaze data and defining the saccades were the same as those described in a previous study^13^. We flagged trials for exclusion when there was no or ill-defined primary saccade, or when the primary saccade showed excessive error (> 5 degree visual angle). We quantified participants’ behavioral memory error as the (signed) difference between the reported location (saccade landing position) and WM target position in polar angle. We included all trials for fMRI data analyses regardless of behavioral exclusion criteria during model estimation to ensure a balanced sampling of spatial positions, but only included trials with reliable behavioral estimates for all subsequent analyses including quantifying the decoding performance and correlating decoded results with behaviors.

### Retinotopic mapping and regions of interest (ROI)

Each participant took part in an 1.5-hour fMRI session for retinotopic mapping. The procedures followed those reported in Mackey et al. ^46^. During the mapping session, participants maintained fixation at the screen center while covertly monitoring a bar aperture swiping through the screen in discrete steps and in four directions: a horizontal bar moving from the top to the bottom, or from bottom to the top, of the screen; a vertical bar moving from the left to the right, or from the right to the left, of the screen. The bar was divided into three rectangular segments (one central segment and two flanking segments) of the same size. Each segment contained a random dot kinematogram (RDK). Participants were required to report whether the RDK in the left (top) or the right (bottom) segment moved in the same direction as the RDK in the central segment by button press before the bar moved into the next step. We adjusted the coherence of the random dot motion to keep the task at about 75% accuracy. Each session contained eight to nine runs. In each run, the bar aperture swept across the screen 12 times, and each sweep consisted of 12 discrete steps. The four sweeping directions were interleaved within each run.

We fit a population receptive field (pRF) model with compressive spatial summation to the BOLD time series of the retinotopic mapping data ^92,93^. To define the ROIs, we visualized the preferred polar angle and eccentricity of the voxels on the cortical surface. We only included voxels whose response variability can be explained by the pRF model over 10%. ROIs were defined by visual inspection by identifying reversals of the voxels’ preferred phase angle on the cortical surface. We defined bilateral dorsal visual ROIs V1, V2, V3, V3AB, IPS0, IPS1, IPS2, IPS3, iPCS and sPCS, each with a full visual field representation.

### MRI acquisition

MRI data were obtained on a Siemens Prisma 3T scanner with a 64-channel head/neck coil. Functional imaging was collected with a voxel size of 2.5^3^ mm and 44 slices (4x simultaneous-multi-slice acceleration; FoV 200 3 200 mm, no in-plane acceleration, TE/TR: 30/750 ms, flip angle: 50 deg, Bandwidth: 2290 Hz/pixel; 0.56 ms echo spacing; P→A phase encoding). The slice prescription was approximately parallel to the calcarine sulcus, covering most of the occipital and parietal lobes, except for the ventral temporal poles and ventral orbitofrontal cortex in some subjects. Spin-echo images in the forward and reverse phase-encoding direction with the same slice prescription and no simultaneous-multi-slice acceleration were intermittently acquired during each scanning session to estimate a field map used to correct for local spatial distortions.

### MRI data preprocessing

T1-weighted anatomical images were segmented and cortical surfaces were constructed using Freesurfer (v6.0). EPI time series data for both the VWM and retinotopic mapping experiments were preprocessed with customized scripts utilizing AFNI functions. We applied B0 field map correction and reverse-polarity phase-encoding (reverse blip) correction on the functional data. All functional data were motion-corrected with 6-parameter affine transform, aligned to the anatomical images, projected onto the cortical surface and re-projected into volume space. This procedure incurred minimal smoothing perpendicular to the cortical surface. Further spatial smoothing was applied only to the retinotopic mapping data using 5 mm FWHM on the cortical surface. Linear trends were removed from the time series. Whenever possible, linear and nonlinear spatial transformations were concatenated into a single transform operation to minimize additional smoothing. For the VWM experiments, the time series was first converted into percentage signal change for each run and then normalized (z-score) across time points within each run.

### Encoding (generative) model

We adapted a generative model based approach to decode the content and the uncertainty of memorized locations (polar angles). This method was first developed by ^41^ and ^44^ to study the neural processes underlying perceptual decision-making when viewing oriented gratings, and was later used by us to investigate the uncertainty of spatial locations maintained in VWM ^13^. Here, we adapted the model to allow for encoding and decoding for two items.

In the generative model, the multivariate voxel pattern given the stimulus location was modeled as a multivariate normal distribution. The mean of the distribution was determined by each voxel’s polar angle tuning function (voxel response as a function of polar angle). The voxel tuning function was modeled by a weighted sum of eight basis functions evenly tiling the location space (Figure 2A). The basis functions are raised sinusoidal functions

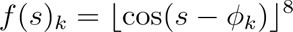

where ⌊⌋ represents half-wave rectification and *ϕ_k_* is the center of the *k*th channel.

We modeled a voxel’s response to two items as a weighted sum of the voxel’s activity for the individual items. Thereby, the response of *i*th voxel *b_i_* given a pair of high- and low-priority items **s** = (*s_H_*, *s_L_*) was modeled as

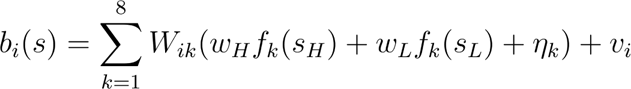

where **W** is a weighting matrix that determines the weights of the basis functions for each voxel, and *w_H_* and *w_L_* are the weights for the high- and low-priority items respectively.

Following the TAFKAP method ^44^, the model considered two sources of noises. First, the noise *η* was specific to each basis function. This noise term was carried over into each voxel through the weighting matrix **W**. It modeled the noise that was shared across voxels with similar tuning functions. *η* followed a zero-mean normal distribution whose covariance matrix was a constant noise magnitude multiplied with an identity matrix *η*~𝒩(0, σ^2^**I**). Second, *v* was the noise specific to each voxel, which followed a zero-mean normal distribution *υ*~𝒩(0, Σ). The covariance matrix Σ was approximated by a rank-one covariance matrix plus a diagonal matrix

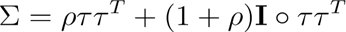

where ° represents the Hadamard product (element-wise product between two matrices), and *τ* is a vector representing the standard deviation of the noise of each voxel. Based on the generative model, the theoretical covariance matrix of the multivariate voxel pattern given the stimuli is

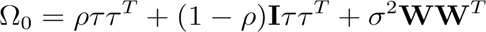

where the first two terms are associated with the voxel-specific noise *υ*, and the last term is associated with *η*^41^.

In addition to the theoretical covariance matrix, the model also considered the empirical sample covariance

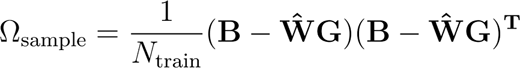

where **B** is the training data and **G** is the response of the basis functions given the training set stimuli. Thus, for each training dataset, we assumed that the voxel activity pattern followed a multivariate normal distribution.

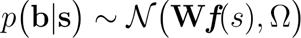

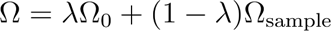

That is, the covariance matrix was estimated as the sample covariance matrix “shrunk” ^94^ to a target covariance matrix, the theoretical covariance matrix Ω_0_. The degree of shrinkage was determined by a free parameter *λ* (see details in ^44^).

### Model fitting and decoding

For each voxel, we averaged the z-scored percentage signal change over a time window late in the delay (5.25 to 12.00 seconds from the delay onset) and treated it as the input to the model. With a passive-viewing experiment, we had previously shown that the decodable voxel activity during this time window is specific to WM and not a mere extension of sensory-evoked response ^13^. In addition, we conducted voxel selection using a previously published dataset, in which the same set of participants performed a similar spatial VWM task but with only one item ^13^. In this 1-item WM experiment, the location of the WM item in each trial was randomly sampled from 32 locations evenly spanning the full circle. We used the averaged voxel activity over the same time window, and applied ANOVA for each voxel using 32 target locations as a categorical independent variable and the voxel response as the dependent variable, and we selected 750 voxels with the highest *F* value for each ROI.

For each participant and each ROI, after selecting the voxels, we used a leave-one-run-out cross-validation procedure to estimate the free parameters in the generative model. The free parameters for the covariance matrix (Ω_sample_, *τ*, *λ*, *ρ* and *σ*) were estimated using the procedures proposed in TAFKAP ^44^. In addition, we fixed the parameter *w_H_* (the weight for the high-priority item) at 1, and estimated a free parameter *w_L_* (the weight for the low-priority item) by a grid search, where we choose the pair of *w_L_* and the weight matrix **W** (estimated by ordinary least squares) that minimizes the mean-square error when estimating the voxel activity of the training data given the stimuli.

After estimating the free parameters using the training data, for each trial in the testset, we decoded a two-dimensional posterior probability distribution of the high- and low-priority items using Bayes rule

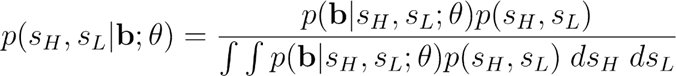

Here *θ* represents the model parameters, and *p*(**b**|*s_H_, s_L_; θ*) is the likelihood function (Figure 2D). The decoder, same as the participants, used the information conveyed by the precue in each trial to perform the task. Specifically, the precue informed the participants the range of the high- and low-priority item in terms of polar angle (Figure 1B). These constraints were implemented as the prior *p*(*s_H_, s_L_*), where *p*(*s_H_, s_L_*) = 1 if *s_H_* ∈ [*a*, *b*] ^ *s_L_* ∈ [*c, d*] and otherwise *p*(*s_H_, s_L_*) = 0 (Figure 2E). Here (*a*, *b*, *c*, *d*) are the boundaries defined by the divider, constraining the range of each of the two items. As polar angle is a circular variable, the boundaries were only defined by two values that are 180° apart. Here we write (*a*, *b*, *c*, *d*) for simplicity.

Lastly, we computed one-dimensional posterior distribution for the high- (or low-) priority item by marginalization

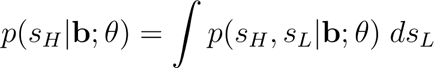

The mean and the standard deviation of this one-dimensional distribution were then used to represent the decoded location and the decoded uncertainty of the high- (or low-) priority item.

### Statistical analysis

To test the effect of prioritization on decoding performance, for each subject and ROI, we computed the magnitude (absolute value) of memory error (reported location minus actual target location) and computed the difference between the high- and low-priority item, and averaged this difference across participants. This difference score was compared to a null distribution obtained by randomly permuting the label (low-priority and high-priority) of each participant’s condition-mean and computing the difference score with the same procedure for 2000 times (Figure 4A). The same procedure was used to test the effect of priority levels on decoded uncertainty (Figure 5A), behavioral memory error (Figure 1C) and saccade response time (Figure 1D).

To investigate whether decoded uncertainty predicts behavioral prioritization on a trial-by-trial basis, for each trial we computed a WM prioritization index as 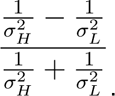 Here, the numerator is the decoding precision (inverse of the square of decoded uncertainty) of the high-priority item minus that of the low-priority item, and the denominator is the sum of the two. For each participant, we then binned the trials into four bins with increasing WM prioritization, and for each bin, we computed a behavioral prioritization index as the magnitude of memory error for the low-priority item minus that of the high-priority item. For each ROI, we pooled the data across all participants (four data points/bins per participant; four bins labeled with four different colors in Figure 5B) after removing the mean from each participant and computed the correlation between neural prioritization and behavioral prioritization. For each ROI, The correlation coefficient was compared to a null distribution estimated by permuting the WM prioritization index and recomputing the correlation coefficient for 2000 times.

We conducted non-parametric bootstrapping to test the significance of the single-trial correlations in several analyses, including the correlation between the magnitude of memory error and response time (Figure 1F), the correlation between response time and the difference of decoded precision between the probed and unprobed items (Figure 6C) and the correlation between response time and the probed item (Supplementary Figure 2). For each ROI, we computed the correlation between the two variables in interest for each participant, and averaged the correlation coefficients across participants. We then resampled the correlation coefficients (with replacement) and computed the averaged correlation coefficients. We repeated this procedure for 2000 iterations to obtain a bootstrapped distribution of the averaged correlation coefficients. The percentage of the iterations in this distribution that was higher or lower than zero was used to compute *p*-values. We reported *p*-values corrected for the number of ROI using FDR with q = 0.05.

### Whole-brain general linear model (GLM)

We conducted a GLM analysis to search for the brain regions with response amplitude that varied with the total WM resource (sum of decoded precision from V3AB over the two items) or neural prioritization index (decoded precision of the high-priority item minus that of the low-priority item in V3AB). The GLM contained the following regressors: (1) stimulus evoked response modeled as a boxcar function with a duration of 500 ms corresponding to the presentation of the WM items. (2) Behavioral response as a boxcar function with a duration of 1500 ms aligned with the response cue (3) The feedback as a boxcar function with a duration of 1000 ms aligned with the onset of the feedback. Lastly, (4) and (5), where the WM delay activity was modeled as two regressors, one as a boxcar function spanning the entire WM delay (12 seconds) with the same amplitude across all trials, and the other with the same timing but with its amplitude modulated trial-by-trial based on the (mean-removed) total WM resource or the (mean-removed) neural prioritization index. In addition, 12 head motion parameters (roll, pitch, yaw, x-, y-, z-translation and their first derivative) estimated during preprocessing were included as nuisance variables to improve the model fits. The above regressors were convolved with a hemodynamic response function modeled as a gamma function which peaked 4.6 seconds after the onset of an event. We conducted the GLMs by the 3dDeconvolve function in AFNI. The two regressors for WM delay, (4) and (5) above, were implemented using the ‘AM2’ option in 3dDeconvolve.

We applied GLM to the BOLD time series (converted to percentage signal change) of each voxel in the volume space. Because we aim to identify the brain regions with their response varying with total WM resource or WM prioritization, we focused on the estimated regression coefficient 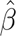 of the fifth regressor above. We mapped this coefficient 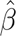 from each participant’s volumetric space to the standard-mesh surface (std.141 space in SUMA) and smoothed each participant’s data on the surface with a targeted fwhm at 5 mm. We conducted group-level statistical tests on the 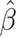 on the cortical surface. We first used 3dttest++ function in AFNI to conduct t-test on each node on the surface. Contiguous nodes that pass this first-level threshold (uncorrected 2-tailed *p* < 0.05) were grouped as clusters. These clusters were deemed significant if their spatial extent passed the size threshold (set at *p* < 0.05) estimated using SurfClust and slow_surf_clustsim function in AFNI.

---

**Supplementary Figure 1.**
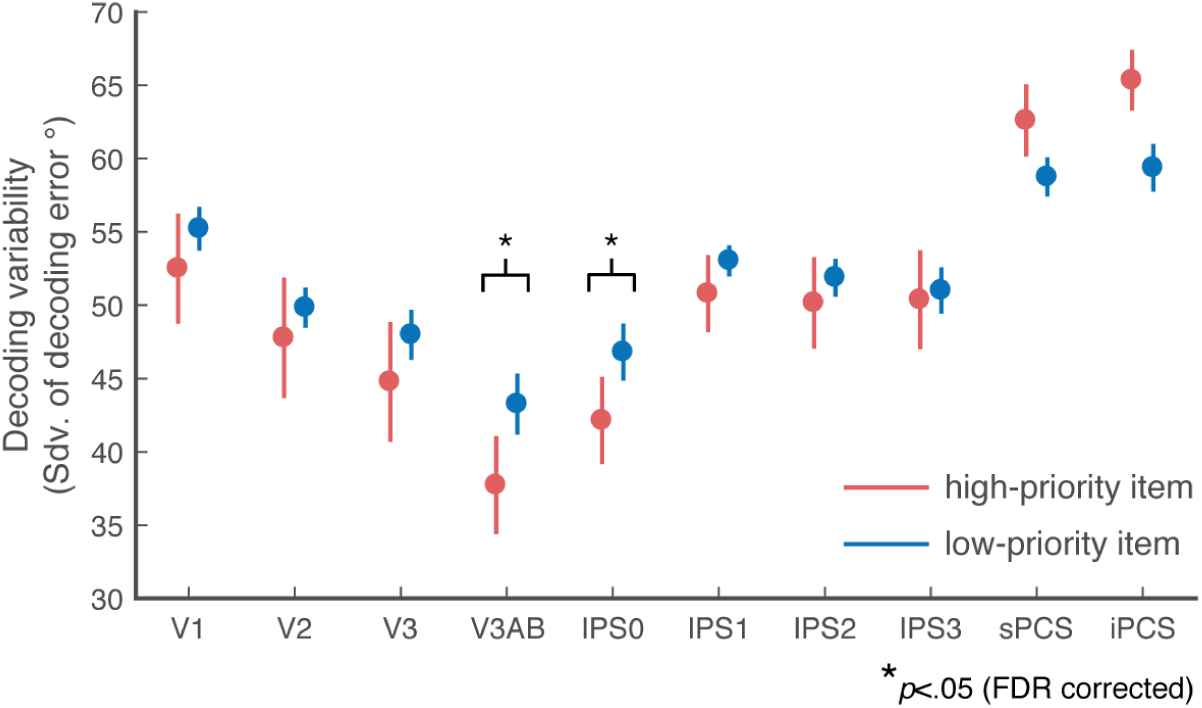
Decoding performance across priority levels. Similar to Figure 4A, but with decoding performance quantified by decoding variability (standard deviation of decoding error) for the high- and low-priority items across ROIs. Data points represent mean ± s.e.m.

**Supplementary Figure 2.**
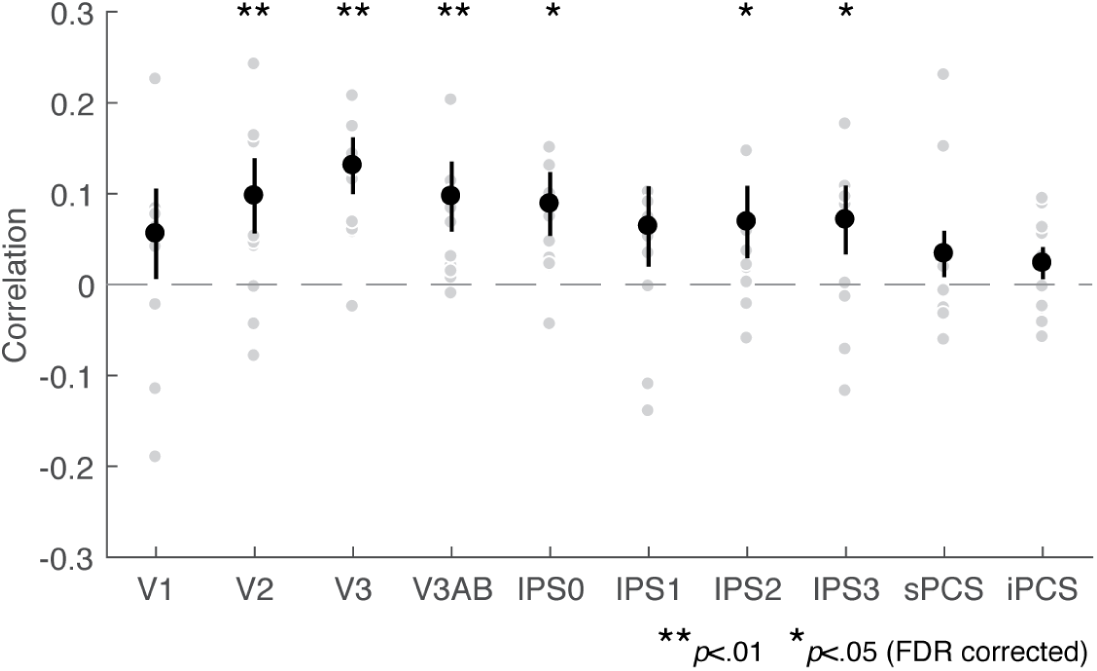
The correlation between saccade response time and the decoded uncertainty of the probed item. Black data points represent mean ± s.e.m. Gray data points represent individual participants.

**Supplementary Figure 3.**
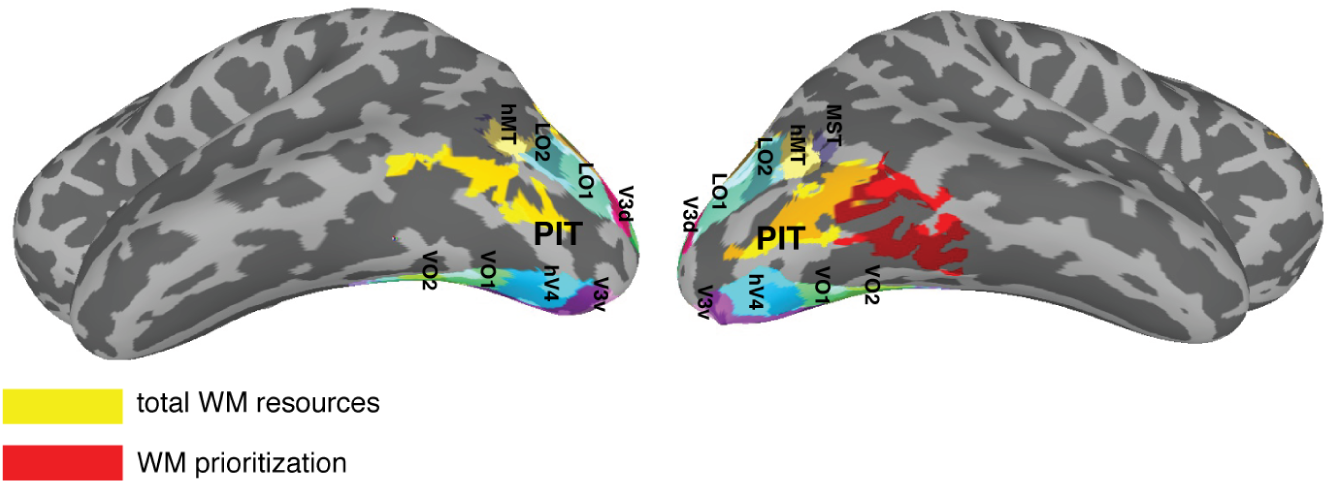
Comparisons between PIT related to WM control with clusters of cortical visual map. PIT is positioned between the lateral visual maps—LO1, LO2, hMT, MSF—and the ventral visual field maps—hV4, VO1, VO2 (the other two ventral maps PHC1 and PHC2 anterior to VO2 can not be seen here). The lateral and the ventral visual field maps shown here are Probability Atlas by Wang et al.^95^.

## Reference

1. Wilken, P. & Ma, W. J. A detection theory account of change detection. J. Vis. 4, 1120–1135 (2004).

2. Alvarez, G. A. & Cavanagh, P. The capacity of visual short-term memory is set both by visual information load and by number of objects. Psychol. Sci. 15, 106–111 (2004).

3. Bays, P. M. & Husain, M. Dynamic shifts of limited working memory resources in human vision. Science 321, 851–854 (2008).

4. Bays, P. M., Catalao, R. F. G. & Husain, M. The precision of visual working memory is set by allocation of a shared resource. J. Vis. 9, 7.1–11 (2009).

5. Adam, K. C. S., Vogel, E. K. & Awh, E. Clear evidence for item limits in visual working memory. Cogn. Psychol. 97, 79–97 (2017).

6. Luck, S. J. & Vogel, E. K. Visual working memory capacity: from psychophysics and neurobiology to individual differences. Trends Cogn. Sci. 17, 391–400 (2013).

7. Rademaker, R. L., Tredway, C. H. & Tong, F. Introspective judgments predict the precision and likelihood of successful maintenance of visual working memory. J. Vis. 12, 21 (2012).

8. Fougnie, D., Suchow, J. W. & Alvarez, G. A. Variability in the quality of visual working memory. Nat. Commun. 3, 1229 (2012).

9. Suchow, J. W., Fougnie, D. & Alvarez, G. A. Looking inward and back: Real-time monitoring of visual working memories. J. Exp. Psychol. Learn. Mem. Cogn. 43, 660–668 (2017).

10. van den Berg, R., Yoo, A. H. & Ma, W. J. Fechner’s law in metacognition: A quantitative model of visual working memory confidence. Psychol. Rev. 124, 197–214 (2017).

11. Honig, M., Ma, W. J. & Fougnie, D. Humans incorporate trial-to-trial working memory uncertainty into rewarded decisions. Proc. Natl. Acad. Sci. U. S. A. 117, 8391–8397 (2020).

12. Yoo, A. H., Acerbi, L. & Ma, W. J. Uncertainty is maintained and used in working memory. J. Vis. 21, 13 (2021).

13. Li, H.-H., Sprague, T. C., Yoo, A. H., Ma, W. J. & Curtis, C. E. Joint representation of working memory and uncertainty in human cortex. Neuron 109, 3699–3712.e6 (2021).

14. Li, A. Y. & Sprague, T. C. Awareness of the relative quality of spatial working memory representations. Atten. Percept. Psychophys. 85, 1710–1721 (2023).

15. Emrich, S. M., Lockhart, H. A. & Al-Aidroos, N. Attention mediates the flexible allocation of visual working memory resources. J. Exp. Psychol. Hum. Percept. Perform. 43, 1454–1465 (2017).

16. Dube, B., Emrich, S. M. & Al-Aidroos, N. More than a filter: Feature-based attention regulates the distribution of visual working memory resources. J. Exp. Psychol. Hum. Percept. Perform. 43, 1843–1854 (2017).

17. Yoo, A. H., Klyszejko, Z., Curtis, C. E. & Ma, W. J. Strategic allocation of working memory resource. Sci. Rep. 8, 16162 (2018).

18. Yoo, A. H. et al. Behavioral Prioritization Enhances Working Memory Precision and Neural Population Gain. J. Cogn. Neurosci. 34, 365–379 (2022).

19. Bays, P. M. Noise in neural populations accounts for errors in working memory. J. Neurosci. 34, 3632–3645 (2014).

20. Klyszejko, Z., Rahmati, M. & Curtis, C. E. Attentional priority determines working memory precision. Vision Res. 105, 70–76 (2014).

21. Brissenden, J. A., Adkins, T. J., Hsu, Y. T. & Lee, T. G. Reward influences the allocation but not the availability of resources in visual working memory. bioRxiv 2021.06.08.447414 (2021) doi:10.1101/2021.06.08.447414.

22. Kastner, S., Pinsk, M. A., De Weerd, P., Desimone, R. & Ungerleider, L. G. Increased activity in human visual cortex during directed attention in the absence of visual stimulation. Neuron 22, 751–761 (1999).

23. Gandhi, S. P., Heeger, D. J. & Boynton, G. M. Spatial attention affects brain activity in human primary visual cortex. Proc. Natl. Acad. Sci. U. S. A. 96, 3314–3319 (1999).

24. Somers, D. C., Dale, A. M., Seiffert, A. E. & Tootell, R. B. Functional MRI reveals spatially specific attentional modulation in human primary visual cortex. Proc. Natl. Acad. Sci. U. S. A. 96, 1663–1668 (1999).

25. Buracas, G. T. & Boynton, G. M. The effect of spatial attention on contrast response functions in human visual cortex. J. Neurosci. 27, 93–97 (2007).

26. Pestilli, F., Carrasco, M., Heeger, D. J. & Gardner, J. L. Attentional enhancement via selection and pooling of early sensory responses in human visual cortex. Neuron 72, 832–846 (2011).

27. Sprague, T. C. & Serences, J. T. Attention modulates spatial priority maps in the human occipital, parietal and frontal cortices. Nat. Neurosci. 16, 1879–1887 (2013).

28. Jerde, T. A., Merriam, E. P., Riggall, A. C., Hedges, J. H. & Curtis, C. E. Prioritized maps of space in human frontoparietal cortex. J. Neurosci. 32, 17382–17390 (2012).

29. Kay, K. N., Weiner, K. S. & Grill-Spector, K. Attention reduces spatial uncertainty in human ventral temporal cortex. Curr. Biol. 25, 595–600 (2015).

30. Sprague, T. C., Ester, E. F. & Serences, J. T. Restoring Latent Visual Working Memory Representations in Human Cortex. Neuron 91, 694–707 (2016).

31. Rahmati, M., Saber, G. T. & Curtis, C. E. Population Dynamics of Early Visual Cortex during Working Memory. J. Cogn. Neurosci. 30, 219–233 (2018).

32. Ester, E. F., Nouri, A. & Rodriguez, L. Retrospective Cues Mitigate Information Loss in Human Cortex during Working Memory Storage. J. Neurosci. 38, 8538–8548 (2018).

33. Sprague, T. C., Itthipuripat, S., Vo, V. A. & Serences, J. T. Dissociable signatures of visual salience and behavioral relevance across attentional priority maps in human cortex. Journal of Neurophysiology vol. 119 2153–2165 Preprint at 10.1152/jn.00059.2018 (2018).

34. Paradiso, M. A. A theory for the use of visual orientation information which exploits the columnar structure of striate cortex. Biol. Cybern. 58, 35–49 (1988).

35. Seung, H. S. & Sompolinsky, H. Simple models for reading neuronal population codes. Proc. Natl. Acad. Sci. U. S. A. 90, 10749–10753 (1993).

36. Foldiak, P. in Computation and Neural Systems (eds. Eeckman, F. H. & Bower, J.) (Springer Science & Business Media, 1993).

37. Sanger, T. D. Probability density estimation for the interpretation of neural population codes. J. Neurophysiol. 76, 2790–2793 (1996).

38. Zemel, R. S., Dayan, P. & Pouget, A. Probabilistic interpretation of population codes. Neural Comput. 10, 403–430 (1998).

39. Jazayeri, M. & Movshon, J. A. Optimal representation of sensory information by neural populations. Nat. Neurosci. 9, 690–696 (2006).

40. Ma, W. J., Beck, J. M., Latham, P. E. & Pouget, A. Bayesian inference with probabilistic population codes. Nat. Neurosci. 9, 1432–1438 (2006).

41. van Bergen, R. S., Ma, W. J., Pratte, M. S. & Jehee, J. F. M. Sensory uncertainty decoded from visual cortex predicts behavior. Nature Neuroscience 18, 1728–1730 (2015).

42. van Bergen, R. S. & Jehee, J. F. M. Probabilistic Representation in Human Visual Cortex Reflects Uncertainty in Serial Decisions. J. Neurosci. 39, 8164–8176 (2019).

43. Walker, E. Y., Cotton, R. J., Ma, W. J. & Tolias, A. S. A neural basis of probabilistic computation in visual cortex. Nat. Neurosci. 23, 122–129 (2020).

44. van Bergen, R. S. & Jehee, J. F. M. TAFKAP: An improved method for probabilistic decoding of cortical activity. bioRxiv 2021.03.04.433946 (2021) doi:10.1101/2021.03.04.433946.

45. Geurts, L. S., Cooke, J. R. H., van Bergen, R. S. & Jehee, J. F. M. Subjective confidence reflects representation of Bayesian probability in cortex. Nature human behaviour 6, 294–305 (2022).

46. Mackey, W. E., Winawer, J. & Curtis, C. E. Visual field map clusters in human frontoparietal cortex. Elife 6, (2017).

47. Rahnev, D. et al. The Confidence Database. Nat Hum Behav 4, 317–325 (2020).

48. Kiani, R. & Shadlen, M. N. Representation of confidence associated with a decision by neurons in the parietal cortex. Science 324, 759–764 (2009).

49. Kiani, R., Corthell, L. & Shadlen, M. N. Choice certainty is informed by both evidence and decision time. Neuron 84, 1329–1342 (2014).

50. Zylberberg, A., Fetsch, C. R. & Shadlen, M. N. The influence of evidence volatility on choice, reaction time and confidence in a perceptual decision. Elife 5, (2016).

51. Ghose, G. M. & Maunsell, J. H. R. Attentional modulation in visual cortex depends on task timing. Nature 419, 616–620 (2002).

52. McAdams, C. J. & Reid, R. C. Attention modulates the responses of simple cells in monkey primary visual cortex. J. Neurosci. 25, 11023–11033 (2005).

53. Treue, S. & Maunsell, J. H. Attentional modulation of visual motion processing in cortical areas MT and MST. Nature 382, 539–541 (1996).

54. Pouget, A., Dayan, P. & Zemel, R. S. Inference and computation with population codes. Annu. Rev. Neurosci. 26, 381–410 (2003).

55. Ma, W. J. & Jazayeri, M. Neural coding of uncertainty and probability. Annu. Rev. Neurosci. 37, 205–220 (2014).

56. Itthipuripat, S., Sprague, T. C. & Serences, J. T. Functional MRI and EEG Index Complementary Attentional Modulations. J. Neurosci. 39, 6162–6179 (2019).

57. Zhou, Y., Curtis, C. E., Sreenivasan, K. K. & Fougnie, D. Common Neural Mechanisms Control Attention and Working Memory. J. Neurosci. 42, 7110–7120 (2022).

58. Ikkai, A. & Curtis, C. E. Cortical activity time locked to the shift and maintenance of spatial attention. Cereb. Cortex 18, 1384–1394 (2008).

59. Reynolds, J. H. & Heeger, D. J. The normalization model of attention. Neuron 61, 168–185 (2009).

60. Yu, Q., Teng, C. & Postle, B. R. Different states of priority recruit different neural representations in visual working memory. PLoS Biol. 18, e3000769 (2020).

61. Ma, W. J., Beck, J. M. & Pouget, A. Spiking networks for Bayesian inference and choice. Curr. Opin. Neurobiol. 18, 217–222 (2008).

62. Denison, R. N., Adler, W. T., Carrasco, M. & Ma, W. J. Humans incorporate attention-dependent uncertainty into perceptual decisions and confidence. Proceedings of the National Academy of Sciences 115, 11090–11095 (2018).

63. Hallenbeck, G., Bolaños, A., Sprague, T. & Curtis, C. Frontal and parietal cortex make distinct contributions to the storage and allocation of resources that support WM. J. Vis. 18, 118–118 (2018).

64. Srimal, R. & Curtis, C. E. Persistent neural activity during the maintenance of spatial position in working memory. Neuroimage 39, 455–468 (2008).

65. Riggall, A. C. & Postle, B. R. The relationship between working memory storage and elevated activity as measured with functional magnetic resonance imaging. J. Neurosci. 32, 12990–12998 (2012).

66. Corbetta, M. & Shulman, G. L. Control of goal-directed and stimulus-driven attention in the brain. Nat. Rev. Neurosci. 3, 201–215 (2002).

67. Silver, M. A. & Kastner, S. Topographic maps in human frontal and parietal cortex. Trends Cogn. Sci. 13, 488–495 (2009).

68. Patel, G. H. et al. Functional evolution of new and expanded attention networks in humans. Proc. Natl. Acad. Sci. U. S. A. 112, 9454–9459 (2015).

69. Alves, P. N., Forkel, S. J., Corbetta, M. & Thiebaut de Schotten, M. The subcortical and neurochemical organization of the ventral and dorsal attention networks. Commun Biol 5, 1343 (2022).

70. Sani, I. et al. The human endogenous attentional control network includes a ventro-temporal cortical node. Nat. Commun. 12, 360 (2021).

71. Curtis, C. E. & Sprague, T. C. Persistent Activity During Working Memory From Front to Back. Front. Neural Circuits 15, 696060 (2021).

72. Christophel, T. B., Klink, P. C., Spitzer, B., Roelfsema, P. R. & Haynes, J.-D. The Distributed Nature of Working Memory. Trends Cogn. Sci. 21, 111–124 (2017).

73. Yellin, D., Berkovich-Ohana, A. & Malach, R. Coupling between pupil fluctuations and resting-state fMRI uncovers a slow build-up of antagonistic responses in the human cortex. Neuroimage 106, 414–427 (2015).

74. Murphy, P. R., O’Connell, R. G., O’Sullivan, M., Robertson, I. H. & Balsters, J. H. Pupil diameter covaries with BOLD activity in human locus coeruleus. Hum. Brain Mapp. 35, 4140–4154 (2014).

75. Reimer, J. et al. Pupil fluctuations track fast switching of cortical states during quiet wakefulness. Neuron 84, 355–362 (2014).

76. Joshi, S. & Gold, J. I. Pupil Size as a Window on Neural Substrates of Cognition. Trends Cogn. Sci. 24, 466–480 (2020).

77. Master, S. L., Li, S. & Curtis, C. E. Trying harder: how cognitive effort sculpts neural representations during working memory. bioRxiv (2023) doi:10.1101/2023.12.07.570686.

78. Burlingham, C. S. et al. Task-related hemodynamic responses in human early visual cortex are modulated by task difficulty and behavioral performance. Elife 11, (2022).

79. Roth, Z. N., Ryoo, M. & Merriam, E. P. Task-related activity in human visual cortex. PLoS Biol. 18, e3000921 (2020).

80. Levin, E. J., Brissenden, J. A., Fengler, A. & Badre, D. Predicted utility modulates working memory fidelity in the brain. Cortex 160, 115–133 (2023).

81. Blanke, O. et al. Location of the human frontal eye field as defined by electrical cortical stimulation: anatomical, functional and electrophysiological characteristics. Neuroreport 11, 1907–1913 (2000).

82. Moore, T. & Armstrong, K. M. Selective gating of visual signals by microstimulation of frontal cortex. Nature 421, 370–373 (2003).

83. Moore, T. & Fallah, M. Microstimulation of the frontal eye field and its effects on covert spatial attention. J. Neurophysiol. 91, 152–162 (2004).

84. Gregoriou, G. G., Gotts, S. J., Zhou, H. & Desimone, R. High-frequency, long-range coupling between prefrontal and visual cortex during attention. Science 324, 1207–1210 (2009).

85. Grace E. Hallenbeck, Nathan Tardiff, Thomas C. Sprague, Clayton E. Curtis. Prioritizing of working memory resources depends on prefrontal cortex. bioRxiv (2024).

86. Stemmann, H. & Freiwald, W. A. Attentive Motion Discrimination Recruits an Area in Inferotemporal Cortex. J. Neurosci. 36, 11918–11928 (2016).

87. Stemmann, H. & Freiwald, W. A. Evidence for an attentional priority map in inferotemporal cortex. Proc. Natl. Acad. Sci. U. S. A. 116, 23797–23805 (2019).

88. Yeo, B. T. T. et al. The organization of the human cerebral cortex estimated by intrinsic functional connectivity. J. Neurophysiol. 106, 1125–1165 (2011).

89. Kay, K. N. & Yeatman, J. D. Bottom-up and top-down computations in word- and face-selective cortex. Elife 6, (2017).

90. Ramezanpour, H. & Fallah, M. The role of temporal cortex in the control of attention. Curr Res Neurobiol 3, 100038 (2022).

91. Wandell, B. A., Dumoulin, S. O. & Brewer, A. A. Visual field maps in human cortex. Neuron 56, 366–383 (2007).

92. Dumoulin, S. O. & Wandell, B. A. Population receptive field estimates in human visual cortex. Neuroimage 39, 647–660 (2008).

93. Kay, K. N., Winawer, J., Mezer, A. & Wandell, B. A. Compressive spatial summation in human visual cortex. J. Neurophysiol. 110, 481–494 (2013).

94. Ledoit, O. & Wolf, M. A well-conditioned estimator for large-dimensional covariance matrices. J. Multivar. Anal. 88, 365–411 (2004).

95. Wang, L., Mruczek, R. E. B., Arcaro, M. J. & Kastner, S. Probabilistic Maps of Visual Topography in Human Cortex. Cereb. Cortex 25, 3911–3931 (2015).

